# Comparison of Deep Learning Approaches for Extreme Low-SNR Image Restoration

**DOI:** 10.64898/2026.01.16.700026

**Authors:** Nasreen Elizabeth Buhn, Sriya Reddy Adunur, Joseph Hamilton, Summer Levis, Guy M. Hagen, Jonathan D. Ventura

## Abstract

**Background:** Live-cell fluorescence microscopy enables the study of dynamic cellular processes. However, fluorescence microscopy can damage cells and disrupt these dynamic processes through photobleaching and phototoxicity. Reducing light exposure mitigates the effects of photobleaching and phototoxicity but results in low signal-to-noise ratio (SNR) images. Deep learning provides a solution for restoring these low-SNR images. However, these deep learning methods require large, representative datasets for training, testing, and benchmarking, as well as substantial GPU memory, particularly for denoising large images.

**Results:** We present a new fluorescence microscopy dataset designed to expand the range of imaging conditions and specimens currently available for evaluating denoising methods. The dataset contains 324 paired high/low-SNR images ranging from four to 282 megapixels across 12 sub-datasets that vary in specimen, objective used, staining type, excitation wavelength, and exposure time. The dataset also includes spinning disk confocal microscopy examples and extreme-noise cases. We evaluated three state-of-the-art deep learning denoising models on the dataset: a supervised transformer-based model, a supervised CNN model, and an unsupervised single image model. We also developed an image stitching method that enables large images to be processed in smaller crops and reconstructed.

**Conclusions:** Our dataset provides a diverse benchmark for evaluating deep learning denoising methods, and our stitching method provides a solution to GPU memory constraints encountered when processing large images. Among the evaluated deep learning models, the supervised transformer-based model had the highest denoising performance but required the longest training time.

## Background

Imaging of live cells and tissues is an essential process that enables scientists to observe dynamic cellular activity. Live imaging is commonly performed using fluorescence microscopy, which allows for the detection and tracking of biological molecules with high sensitivity and specificity [1]. These biological molecules are probed with fluorophores, which are excited by distinct wavelengths of light to produce emissions used to generate images. However, during the excitation of fluorophores, photobleaching and phototoxicity can occur, which introduce challenges to the consistency and reproducibility of imaging data [2].

The excitation of fluorophores can result in damage to their chemical structure, a process known as photobleaching. As fluorophores undergo photobleaching, they may interact with oxygen, generating reactive oxygen species (ROS) [2]. An unnatural increase in ROS can cause phototoxicity, leading to detrimental changes in a specimen, including damage to DNA, induced mutations, oxidized proteins, and potential disruption of the developmental processes within a cell [2].

To produce reliable data, phototoxicity must be minimized. Accordingly, numerous approaches have been developed to mitigate its effects. Many strategies focus on reducing a sample’s light exposure by modifying microscope hardware, sample environment, and imaging conditions. One approach has been limiting light exposure to areas outside the focal plane. This approach serves as the basis for multiple fluorescence microscopy modalities, including total internal reflection fluorescence, light sheet fluorescence microscopy, and two-photon microscopy [3]. Despite these methods, phototoxicity remains a challenge, since high local light intensities can still damage cells, especially during prolonged live imaging [2].

Reducing excitation light intensity and/or exposure times can minimize photobleaching and phototoxicity but leads to low signal-to-noise ratio (SNR) images. To circumvent this, computational image restoration techniques have been applied to restore low SNR images. However, traditional restoration algorithms, such as BM3D, which rely on predefined mathematical models and heuristics, struggle to address complex noise patterns [4]. In contrast, deep learning models can learn the structure of complex noise types, including cases with unknown noise levels [5]. Recent work by Hagen et al. demonstrates that these deep learning models outperform traditional models in terms of quantitative metrics and visual quality [6].

Deep learning is a subset of machine learning that uses artificial neural networks, computational networks inspired by the brain, to learn patterns from large datasets. These networks rely on layers of interconnected nodes to transform input data into outputs using learned weights. Among deep learning methods, convolutional neural networks (CNNs) [7] have been proven effective for image-related tasks. CNNs use filters to extract image features like edges, textures, and shapes, allowing the network to learn visual patterns. More recently, transformer-based models, initially developed for natural language processing [8], have been adapted for vision tasks [9], offering advantages in long-range dependencies and global contexts compared to CNNs.

Deep learning approaches for image restoration can be classified into three major categories: supervised, semi-supervised and unsupervised [10]. Supervised methods rely on large datasets of paired low and high SNR images to learn the relationship between noisy and clean images. In contrast, unsupervised methods do not rely on ground truth data. Unsupervised methods include single-image methods, which denoise a single image without prior knowledge of the noise distribution, and self-supervised methods, which rely solely on low SNR images. Semi-supervised methods sit conceptually between supervised and unsupervised methods, relying on both labeled and unlabeled data [9]. Extensive and representative datasets are required to train supervised models and to benchmark the performance of both supervised and unsupervised models.

To meet the data requirements necessary for denoising models, datasets should span fluorescence microscopy imaging modalities and biological specimens. Such datasets should capture complex noise patterns arising from varying imaging conditions and specimens, enabling supervised models to learn robust denoising mappings. This range of conditions allows for the assessment of model generalizability and helps prevent overfitting to specific training data. These datasets serve as valuable benchmarks for comparing deep learning methods for fluorescence microscopy image denoising.

While a variety of datasets are important for fluorescence microscopy denoising, few are currently available. One such dataset, produced by Zhang et al. [12], consists of 12,000 images spanning fluorescence microscopy modalities, including confocal, two-photon, and widefield. This dataset includes samples from cells, zebrafish, and mouse brain tissue, but is limited in both sample diversity and image quality. The W2S dataset produced by Zhou et al. [13], provides 360 image sets with varying noise levels but is constrained to solely widefield imaging. Weigert et al. [14] provide a dataset of paired image patches of Planaria to evaluate denoising methods. More recently, Hagen et al. [6] produced a dataset consisting of 567 paired images ranging from 0.26 to 4.19 megapixels, covering both widefield and confocal imaging modalities. This dataset includes images of actin, mitochondria, nucleus, and membrane samples. However, it remains limited in terms of noise level variation, image sizes, and sample types. A well-suited dataset for training deep learning models should include a diverse range of fluorescence microscopy modalities, objective lenses, noise levels, exposure times, and biological samples.

We introduce a novel dataset of 324 high-resolution images designed to increase the diversity of specimens and imaging modalities represented in existing fluorescence microscopy denoising datasets. Previously unrepresented specimens include breast cancer tissue array spots, earthworm, freshwater fish gills, rat testes, rabbit testes, human ovary, and tubulin. To expand the range of imaging modalities, we include spinning disk confocal microscopy, and extreme-noise cases obtained by widening the gap between high and low exposure times. This increased diversity of specimens, imaging modalities, and noise conditions makes our dataset a more comprehensive and well-suited resource for evaluating the performance of fluorescence microscopy denoising models.

In addition to the necessity of large, diverse datasets, deep learning approaches require substantial memory for computation. As a result, denoising of fluorescence microscopy images is limited by GPU memory constraints. To address this challenge, we introduce an image stitching approach that enables large images to be denoised in smaller crops and reassembled.

We evaluated the performance of three state-of-the-art deep learning denoising models using our dataset. Our selection included the most competitive supervised denoising models [14], [15] as well as a leading unsupervised single-image denoising model [16]. BM3D was excluded from our analysis due to its inferior performance relative to the supervised model CARE in previous work [6].

The three evaluated state-of-the-art deep learning models included: Noise2Fast, Restormer, and CARE. Noise2Fast, proposed by Lequyer et al. [16], is a blind single-shot unsupervised denoiser tailored for speed. Noise2Fast relies on a unique checkerboard-down sampling technique. Checkerboard down-sampling removes half of the pixels present in the original image and rearranges the remaining pixels to fill the gaps. This process generates a set of four images, which are used to train a feed-forward neural network until convergence with the original image is obtained. Content Aware Image Restoration (CARE), developed by Weigert et al. [14], is a supervised model that utilizes a U-Net architecture [17] to learn mappings from degraded images to denoised versions. Lastly, Restormer, introduced by Zamir et al. [15], is a supervised denoising model that employs an encoder-decoder transformer-based architecture. The model’s core components include a Multi-Dconv Head Transposed Attention (MDTA) block and a Gated Dconv Feed-Forward Network (GDFN). MDTA aids the model in learning both fine details and broader patterns by combining attention mechanisms and depth-wise convolutions. GDFN improves feature quality by using gates and convolutional layers to refine and enhance each layer’s image content. The model utilizes a progressive learning approach that begins with smaller image patches and larger batch sizes and gradually shifts to larger image patches and smaller batches.

### Data Description

We generated fifteen distinct datasets of paired low and high-SNR images composed of specimens from actin, breast cancer array, earthworm, fish gill, rat testes, immature ovaries, human ovaries in active phase, mitochondria, mouse brain, rabbit testes, and tubulin. The images range in size from 4.19 to 282.22 megapixels (MP). Imaging conditions varied in light intensity, exposure times, objectives, excitation wavelengths, and stains as detailed in Table 1.

**Table 1.** Overview of dataset imaging conditions.

| Dataset | Number of image pairs | Original image sizes (MP) | Exposure time (low, high, ms) | Objective Mag/NA | Stain | Excitation (nm) | Sample source | Catalog number |
| --- | --- | --- | --- | --- | --- | --- | --- | --- |
| Actin 60X | 23 | 5 - 55 | 1, 100 | 60×/1.42 | Alexa488 | 470 | Invitrogen | Fluoroslides #1 |
| Breast Cancer Array 60X | 1 | 282 | 1, 100 | 60×/1.42 | H&E | 530 | Tissuearray | BR249 |
| Earthworm 20X Super Extreme | 1 | 134 | 1, 600 | 20×/0.45 | H&E | 530 | Eisco | BS18225 |
| Fish Gill 20X | 1 | 256 | 1, 100 | 20×/0.45 | H&E | 530 | Eisco | BS18101 |
| Extreme Fish Gill 10x | 12 | 19 - 68 | 1, 300 | 10×/0.40 | H&E | 530 | Eisco | BS18101 |
| Spinning Disk Rat Testis 60X | 105 | 4 | 20, 200 | 60×/1.35 | H&E | 532 | Carolina | 316464 |
| Spinning Disk Rat Testis 60X Medium Extreme | 100 | 4 | 20, 500 | 60 ×/1.35 | H&E | 450 | Carolina | 316464 |
| Fish Gill 10X Super Extreme | 11 | 28 - 55 | 1, 600 | 10×/0.40 | H&E | 530 | Eisco | BS18101 |
| Human Ovary Active Phase 10X | 13 | 28 - 72 | 1, 100 | 10×/0.4 | H&E | 530 | Carolina | 616024 |
| Human Ovary Active Phase 4X | 11 | 28 - 55 | 1, 100 | 4×/0.16 | H&E | 530 | Carolina | 616024 |
| Immature Ovary 20X Super Extreme | 1 | 90 | 1, 600 | 20×/0.45 | H&E | 530 | Eisco | BS18222 |
| Mitochondria 60X | 14 | 5 - 55 | 1, 100 | 60×/1.42 | Mito Tracker | 530 | Invitrogen | Fluoroslides #1 |
| Mouse Brain 10X | 8 | 37 - 126 | 1, 100 | 10×/0.4 | GFP | 470 | Sunjin Lab | - |
| Rabbit Testis 10X | 9 | 5 - 81 | 1, 100 | 10×/0.40 | H&E | 530 | Amscope | PS50 |
| Tubulin 60X | 14 | 5 - 55 | 1, 100 | 60×/1.42 | BODIPY | 470 | Invitrogen | Fluoroslides #2 |

### Analyses

Adaptive stitching refers to stitching performed with intensity adjustment of 640 x 640 overlapping test crops. Non-adaptive stitching refers to the assembly of 512 x 512 non-overlapping test crops into a composite image without adjustment.

The denoising performance of Restormer, Noise2Fast, and CARE is summarized in Table 2 and Figures 1-3. Figures 1 and 2 visually present average SSIM and PSNR results for each model across all test datasets. Table 2 presents these metrics numerically. Figure 3 shows representative low-SNR input images, denoised outputs from each model, and the corresponding high-SNR ground truth images from each dataset. Restormer consistently achieved the highest average PSNR and SSIM, as shown in Table 2 and visualized in Figures 1 and 2. CARE and Noise2Fast’s comparative performance varied depending on the datasets, as shown in Table 2 and Figures 1 and 2.

**Table 2.** Average PSNR and SSIM denoising results. Test image sets, consisting of the last 10% of image pairs from each dataset, were denoised by each model following training. PSNR and SSIM values were averaged across the datasets. The highest PSNR and SSIM values per dataset are bolded.

| Dataset | PSNR |  |  |  | SSIM |  |  |  |
| --- | --- | --- | --- | --- | --- | --- | --- | --- |
|  | Raw | Care | Restormer | Noise2Fast | Raw | Care | Restormer | Noise2Fast |
| Actin 60X | 23.22 | 29.5 | **31.01** | 29.47 | 0.37 | 0.65 | **0.68** | 0.65 |
| Breast Cancer Array 60X | 23.31 | 28.07 | **34.26** | 30.15 | 0.4 | 0.74 | **0.82** | 0.78 |
| Earthworm 20X Super Extreme | 17.15 | 18.54 | **19.89** | 18.62 | 0.08 | 0.12 | **0.18** | 0.15 |
| Fish Gill 20X | 22.32 | 32.04 | **33.41** | 32.38 | 0.39 | 0.85 | **0.87** | 0.85 |
| Extreme Fish Gill 10x | 15.7 | 20.83 | **23.90** | 20.02 | 0.05 | 0.28 | **0.43** | 0.23 |
| Spinning Disk Rat Testis 60X | 27.66 | 30.28 | **33.73** | 29.37 | 0.75 | 0.89 | **0.93** | 0.88 |
| Spinning Disk Rat Testis 60X Medium Extreme | 18.09 | 26.6 | **27.64** | 26.6 | 0.14 | 0.49 | **0.53** | 0.49 |
| Fish Gill 10X Super Extreme | 15.4 | 20.33 | **23.16** | 19.14 | 0.05 | 0.28 | **0.42** | 0.2 |
| Human Ovary Active Phase 10X | 19.78 | 25.88 | **26.80** | 25.94 | 0.34 | 0.64 | **0.68** | 0.65 |
| Human Ovary Active Phase 4X | 21.14 | 27.03 | **28.12** | 26.83 | 0.43 | 0.68 | **0.72** | 0.66 |
| Immature Ovary 20X Super Extreme | 14.12 | 18.06 | **21.43** | 17.41 | 0.03 | 0.22 | **0.39** | 0.16 |
| Mitochondria 60X | 22.86 | 30.38 | **32.16** | 30.44 | 0.3 | 0.7 | **0.75** | 0.69 |
| **Mouse Brain 10X** | 18.35 | 24.43 | **28.24** | 25.71 | 0.11 | 0.37 | **0.50** | 0.42 |
| **Rabbit Testis 10X** | 21.21 | 29.35 | **30.60** | 29.24 | 0.39 | 0.77 | **0.79** | 0.76 |
| **Tubulin 60X** | 18.56 | 27.15 | **28.73** | 27.07 | 0.16 | 0.6 | **0.66** | 0.57 |

**Figure 1.**
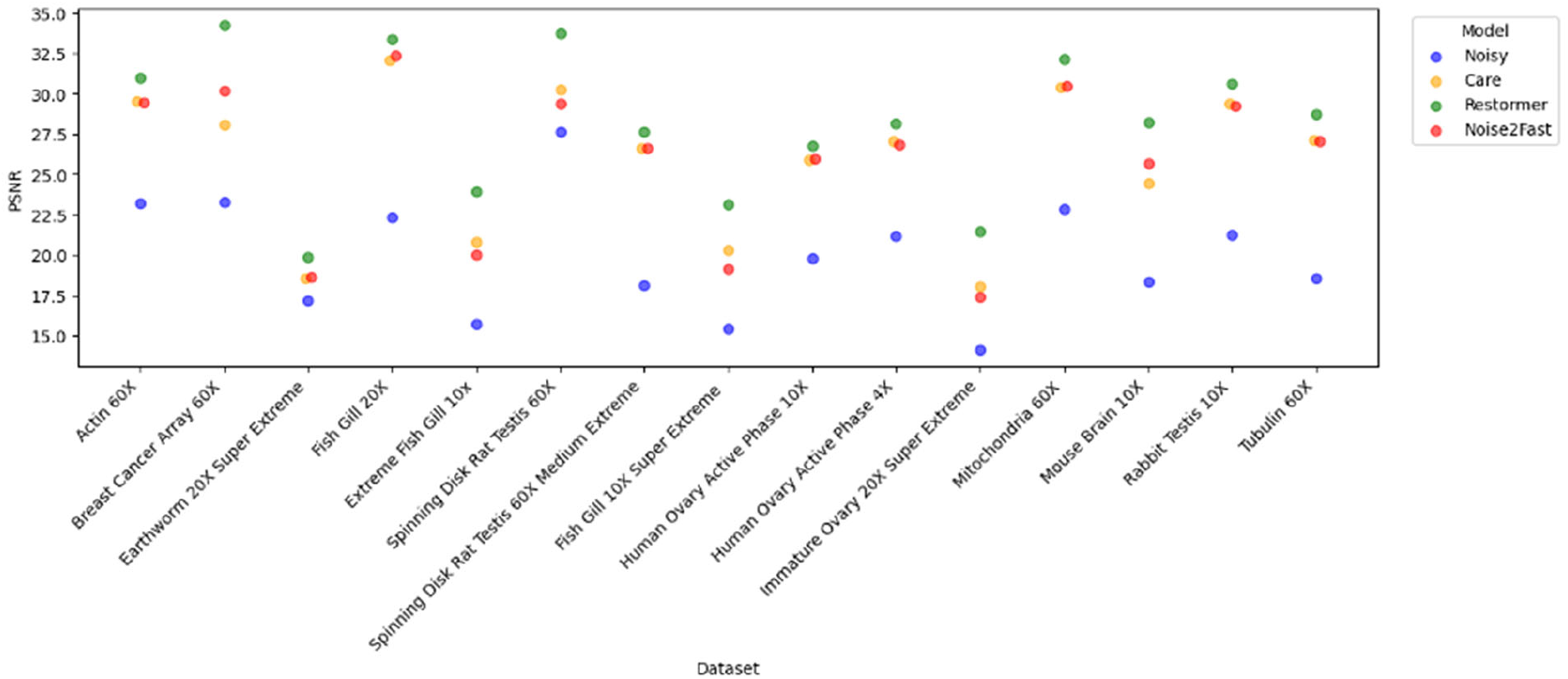
Restormer achieves the highest PSNR on all test datasets. PSNR results from Table 2 are represented visually; models with similar results for a given dataset are horizontally jittered.

**Figure 2.**
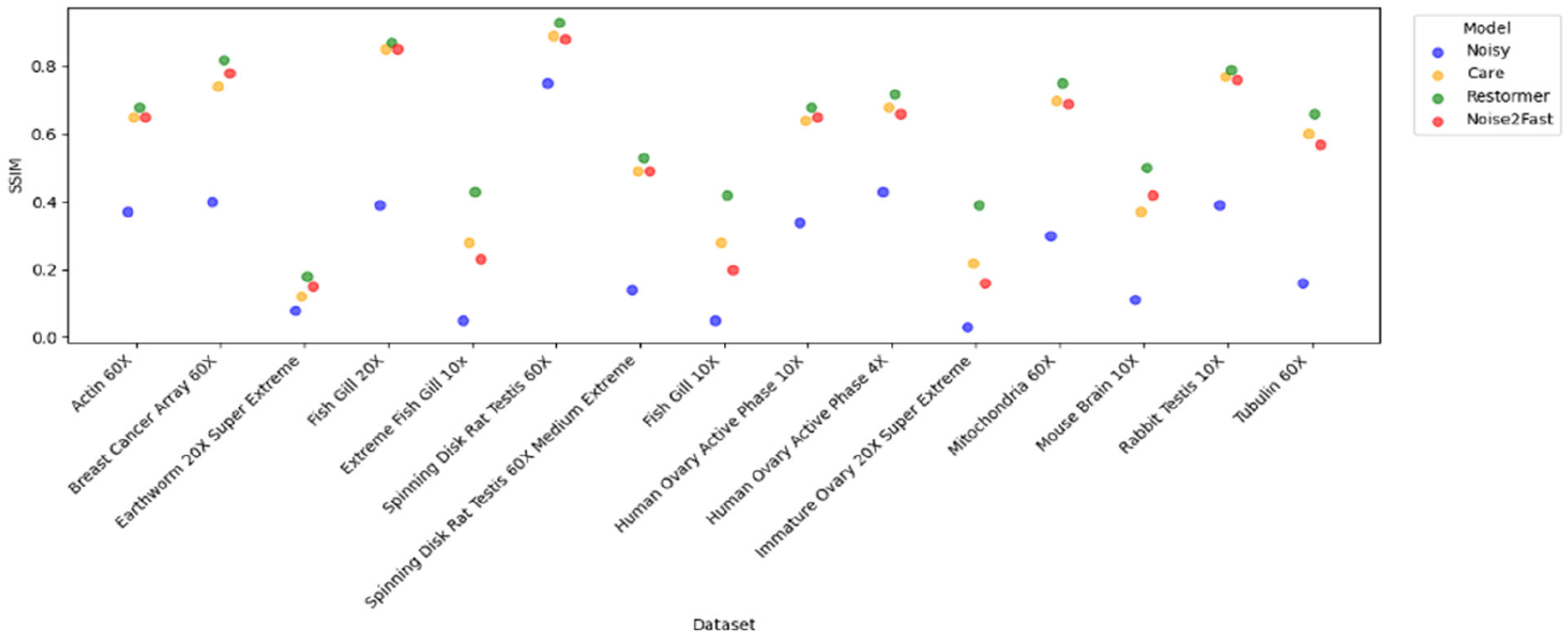
Restormer achieves the highest SSIM for all test datasets. SSIM results from Table 2 are represented visually; models with similar results for a given dataset are horizontally jittered.

**Figure 3.**
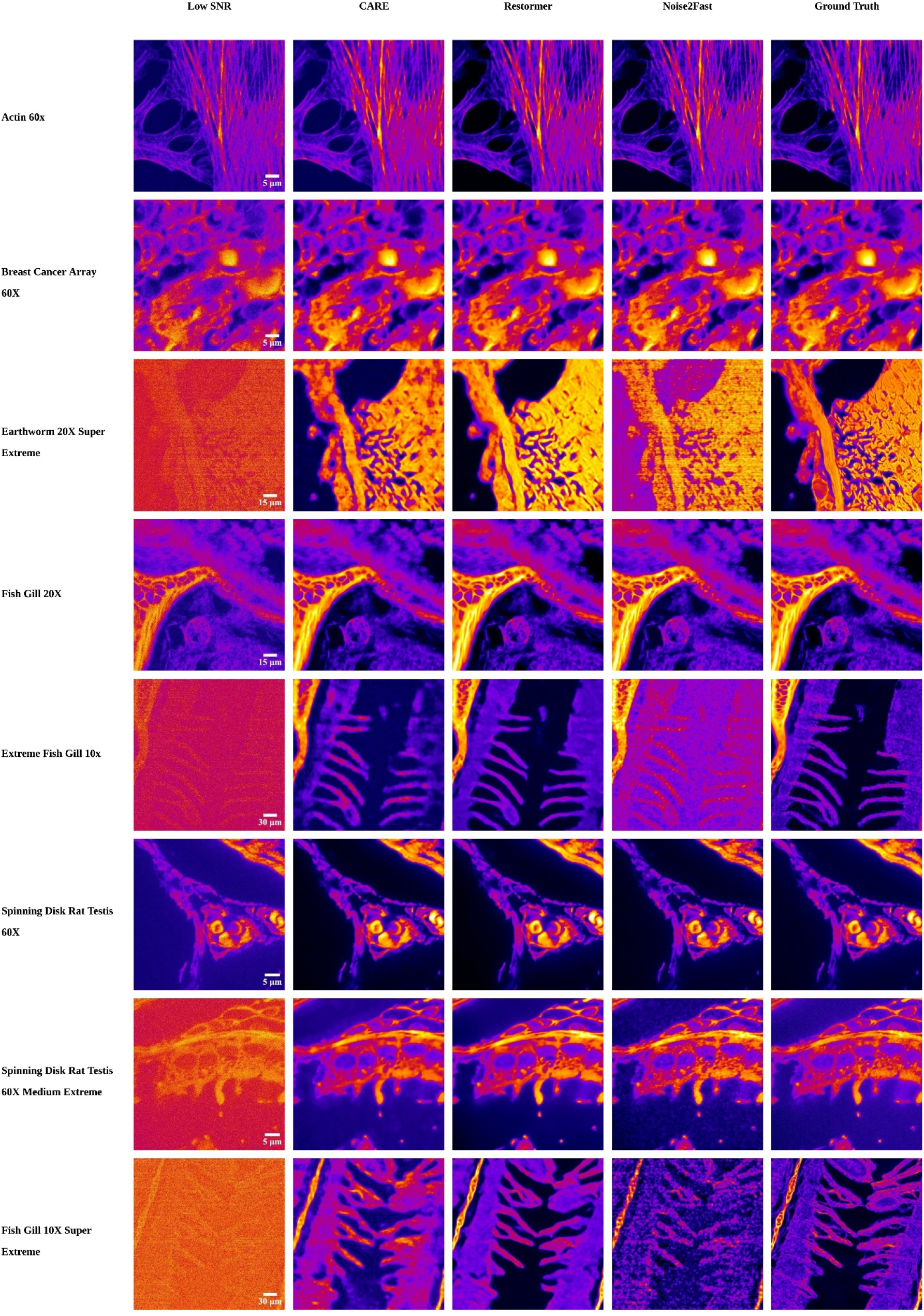
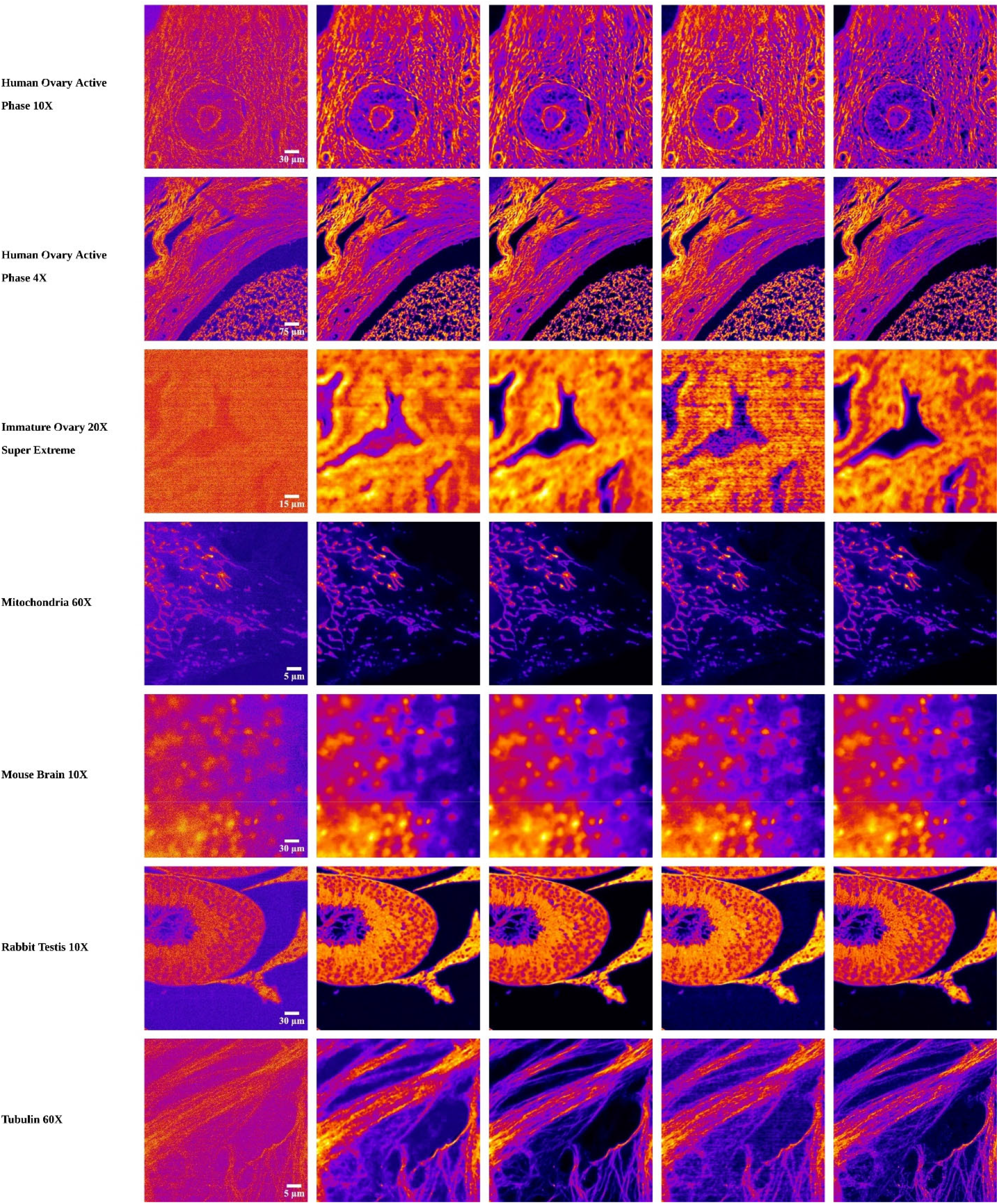
Example denoising results for each model across all test datasets. A single representative image from each test dataset is shown under low SNR conditions, followed by the denoised outputs of CARE, Restormer, and Noise2Fast models and corresponding ground truth image. Denoised outputs were generated by applying the corresponding trained model to the respective low SNR images.

Restormer’s training times ranged from 16-18 hours to several days, while the CARE network trained in approximately 3.5 hours per dataset. Noise2Fast does not require training on a separate training set. Inference times also varied between models: Restormer required approximately 0.45 seconds per image, CARE required about 1 second per image, and Noise2Fast, due to its iterative self-supervised denoising method, took between 9 and 40 seconds per image.

The deep learning models introduced variations in light intensity between neighboring denoised crops, as shown in Figure 4. Our adaptive image stitching method reduces these intensity differences across stitched crops compared to non-adaptive stitching, as shown in Figure 4. Figure 5 shows an example adaptively stitched image compared to its corresponding low and high SNR images.

**Figure 4.**
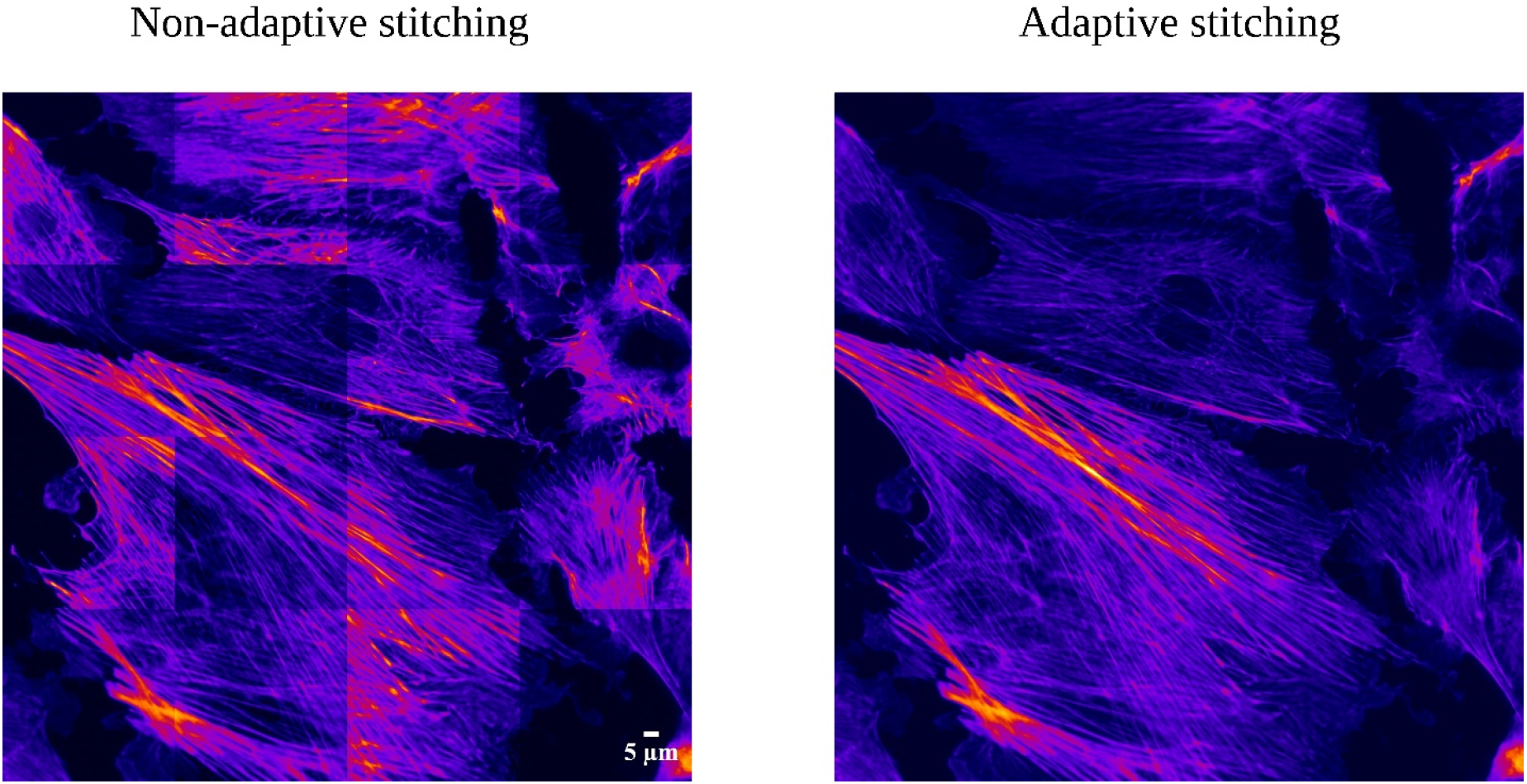
Effect of stitching logic on Actin-60x denoised images. The left panel shows non-adaptively stitched 512×512 non-overlapping crops. The right panel shows adaptively stitched 640x640 overlapping crops stitched using intensity-adjustment stitching logic. Final images are 2048×2048 pixels.

**Figure 5.**
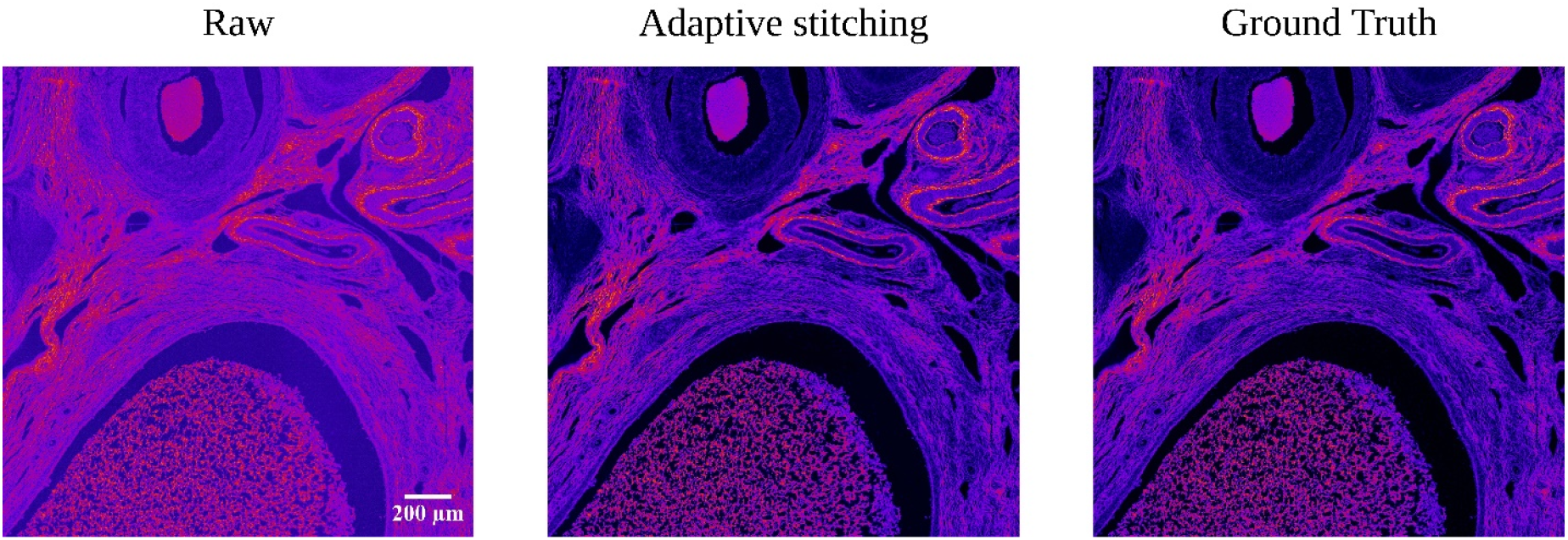
Example adaptively stitched image from the human ovary active phase 10x dataset. The low-SNR image is shown to the left, followed by the adaptively stitched reconstruction containing 36 individually denoised overlapping crops, and the corresponding ground truth image. All images are 3072×3072 pixels.

## Discussion

1. Across the specimens, imaging modalities, and noise conditions present in our dataset, Restormer achieved the highest performance in terms of PSNR and SSIM. However, due to its supervised training and Transformer-based encoder-decoder architecture, Restormer required substantially longer training times than the supervised, U-net based CARE model and self-supervised Noise2Fast model.

While its computational costs were considerably higher, Restormer was the most effective model for removing noise while preserving the structural integrity of the original images. Preserving the fine structural detail of fluorescence microscopy images is particularly important because the introduction of artifacts compromises the reliability of downstream biological analysis. This highlights the critical trade-off between computational cost and model accuracy in fluorescence microscopy denoising.

In addition to computational requirements, supervised deep learning models rely on representative, paired image datasets to learn robust mappings from low-SNR to high-SNR images. Denoising performance depends on a model’s ability to learn these mappings, but with limited or unrepresentative training datasets, models risk overfitting to specific imaging conditions and fail to handle the inherent variability of live fluorescence microscopy imaging. In contrast, unsupervised approaches do not require extensive training data, but they demonstrated inferior denoising performance, as they struggled to remove noise and maintain the structural integrity of the images in our dataset.

The deep learning models introduced variations in light intensity between the individually denoised neighboring crops, resulting in visible seams and uneven lighting in the reconstructed images. To address the issue of uneven lighting in denoised crops, we implemented an adaptive stitching approach that adjusts each image segment’s light intensity based on its neighbors. This method was able to effectively decrease intensity variation across images. Our adaptive stitching process enabled the denoising of large images using deep learning models without exceeding our memory constraints.

### Potential Implications

Our results highlight the relationship between computational costs and denoising accuracy, underscoring the need for deep learning denoising models that are both computationally efficient and able to remove noise without compromising image integrity. Additionally, given the diversity of biological structures included in our dataset, it would be valuable to explore whether certain deep learning techniques are more effective for denoising specific biological structures and how different stains may impact denoising accuracy. Lastly, since tiny structural distortions can have a significant impact on biological interpretations but may not strongly affect PSNR and SSIM, there is a need for image quality metrics specifically tailored to microscopy images [18-20].

## Methods

### Microscopy

We acquired the fluorescence images using an Olympus BX53 microscope equipped with a motorized XY stage (Applied Scientific Imaging, Eugene, OR), Aura III light source (Lumencor, Beaverton, OR), Fluorescence filters (Chroma, Bellows Falls, VT), and Fusion BT camera (Hamamatsu Photonics, Hamamatsu, Japan). The spinning disk confocal setup is described in our previous work [24].

### Deep learning methods

Images were segmented into 512×512 non-overlapping crops, resulting in a collection of 17,568 total crops. Each image pair dataset was split into training, validation, and testing sets. Ninety percent of each dataset was allocated for training, from which five image pairs were reserved for validation. The remaining holdout, ten percent of each dataset was used for testing. Each model was trained and tested using an Nvidia V100 32GB GPU.

Noise2Fast was trained according to the authors’ implementation with a sequence of nine 3×3 convolutions using ReLU activation followed by a sigmoid output activation [21]. The model was optimized using a binary cross-entropy loss function and the Adam optimizer, with a learning rate of 1 × 10^-3^ and a batch size of 1. No batch normalization was applied.

The CARE model was implemented using the CSBdeep framework, and its convolutional layers were optimized with a mean absolute error (MAE) loss function, as developed by the authors [22]. Training was performed for 400 epochs using a batch size of 16 and a patch size of 128×128.

The Restormer model was trained using the AdamW optimizer and L1 loss. The initial learning rate was 3 × 10^-3^, which was reduced to 1 × 10^-6^ over 300,000 iterations. Training started with an initial patch size of 128 x 128 and batch size of 64 and ended with a final patch size of 384x384 and batch size of 8.

Training followed the specifications of Zamir et al. with the following modifications: a single GPU was used with 1 input and 1 output channel [18]. Images were processed as 512×512 non-overlapping crops. The crops were filtered to regions containing substantial non-background signal, normalized using percentile-based normalization with percentiles determined from the corresponding raw images, and cropped identically in paired low and high-SNR images.

### Image quality metrics

We analyzed the performance of the models on a holdout test set from each dataset not used during training. Performance was quantified using peak signal-to-noise ratio (PSNR) and structural similarity index measure (SSIM) by comparing each denoised result to its corresponding ground truth image. PSNR measures the ratio of the maximum signal power in an image to the power of the noise present in the image. SSIM is based on human visual perception of an image and evaluates image similarity using contrast, luminance, and structure. SSIM values range from zero to one. Higher PSNR and SSIM values indicate increased image similarity and, thus, greater denoising success.

Prior to calculating PSNR and SSIM, images were normalized as described in Supplementary Notes, Section 2.2 of Weigert et al. [12]. Percentile-based normalization was applied to each high-SNR image using the 0.1st and 99.9th percentiles. The low-SNR, high-SNR, and denoised images were then mean-centered. The low-SNR and denoised images were scaled by a factor equal to their covariance with the high-SNR image, divided by their own variance.

### Adaptive image stitching

We prepared the full-size test images for stitching by adding 64-pixel reflective padding and dividing them into overlapping crops. Each image was padded, then segmented into 640×640-pixel crops, each overlapping with its neighboring tile or the padding by 64 pixels, as shown in Figure 6. The central region of each crop, excluding the overlap, represents the valid region. The valid region measured 512×512-pixels. Crops were saved with their coordinates for relocation.

**Figure 6.**
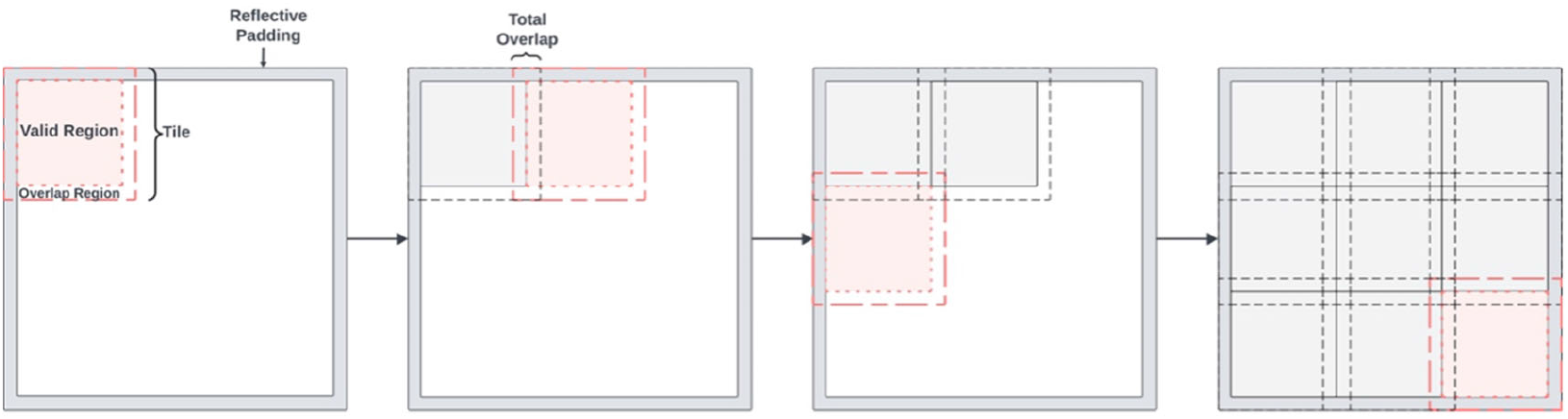
Image cropping logic. The red tile shows the progression of the cropping logic. The shaded area marks the valid region of each tile, while the overlapping region is the space between the valid region and the outer dashed line. Cropping logic proceeds until the entire image is processed.

After cropping, we denoised, intensity-corrected, weight-adjusted, and aligned crops to produce uniform composite images. Crops were denoised using the trained Restormer model, then passed to an adaptive stitching algorithm that identified the overlapping regions between the current tile and its neighbors. The trust region reflective algorithm over a least squares objective was used to optimize the agreement between overlapping regions by adjusting each tile’s scale and shift relative to its neighbors. To anchor the adaptive stitching algorithm, we identified the undenoised tile with the greatest intensity and assigned its corresponding tile in the denoised image a fixed scale and shift. Following intensity adjustment, a feathered weight mask was applied to each tile, with weights linearly decreasing from the center of each tile to its edges. In overlapping regions, pixels from all neighboring crops were summed and divided by the total weight at each position, producing smooth transitions between crops.

## Availability of source code and requirements

Project name: Comparison of Deep Learning Approaches for Extreme Low-SNR Image Restoration. https://github.com/nazbuhn/Comparison-of-Deep-Learning-Approaches-for-Extreme-Low-SNR-Image-Restoration

Operating systems(s): Platform independent. Programming language(s): Python, MATLAB

### Abbreviations

BM3D: block matching and 3D filtering
CARE: content-aware image restoration
CNN: convolutional neural network
GDFN: Gated-Dcov Feed-Forward Network
GPU: graphics processing unit
MDTA: multi-Dconv transposed attention
MP: megapixels
NA: numerical aperture
PSNR: peak signal-to-noise ratio
ROS: reactive oxygen species
SNR: signal to noise ratio
SSIM: structural similarity index measure.

## Author Contributions

NEB: analyzed data, wrote the paper; SRA: analyzed data, wrote the paper; JH: acquired data; SL: acquired data; JV: conceived project, analyzed data, supervised research, wrote the paper; GH: conceived project, acquired data, analyzed data, supervised research, wrote the paper.

